# Coalescent-based species delimitation meets deep learning: Insights from a highly fragmented cactus system

**DOI:** 10.1101/2020.12.23.424219

**Authors:** Manolo F. Perez, Isabel A. S. Bonatelli, Monique Romeiro-Brito, Fernando F. Franco, Nigel P. Taylor, Daniela C. Zappi, Evandro M. Moraes

## Abstract

Delimiting species boundaries is a major goal in evolutionary biology. An increasing body of literature has focused on the challenges of investigating cryptic diversity within complex evolutionary scenarios of speciation, including gene flow and demographic fluctuations. New methods based on model selection, such as approximate Bayesian computation, approximate likelihoods, and machine learning are promising tools arising in this field. Here, we introduce a framework for species delimitation using the multispecies coalescent model coupled with a deep learning algorithm based on convolutional neural networks (CNNs). We compared this strategy with a similar ABC approach. We applied both methods to test species boundary hypotheses based on current and previous taxonomic delimitations as well as genetic data (sequences from 41 loci) in *Pilosocereus aurisetus*, a cactus species complex with a sky-island distribution and taxonomic uncertainty. To validate our method, we also applied the same strategy on data from widely accepted species from the genus *Drosophila*. The results show that our CNN approach has high capacity to distinguish among the simulated species delimitation scenarios, with higher accuracy than ABC. For the cactus dataset, a splitter hypothesis without gene flow showed the highest probability in both CNN and ABC approaches, a result agreeing with previous taxonomic classifications and in line with the sky-island distribution and low dispersal features of *P. aurisetus*. Our results highlight the cryptic diversity within the *P. aurisetus* complex and show that CNNs are a promising approach for distinguishing complex evolutionary histories, even outperforming the accuracy of other model-based approaches such as ABC.

## Introduction

Recognizing species boundaries has long been a major challenge for biologists. The major difficulty is to some degree related to the numerous existing species concepts. The use of specific definitions can lead to alternative strategies for identifying species boundaries in empirical datasets (de Queiroz 2007; Carstens et al. 2013). However, different species concepts can be considered elements of diverse properties that are associated with the dynamics of the speciation continuum. After the proposal of the unified species concept (Queiroz, 2007), the roles of species concept theory and species delimitation methodologies became apparent. The view of species as independent segments of a metapopulation indicated lineage independence as the only criterion necessary for delimiting species boundaries, avoiding any disagreement purely related to the species concept.

Selecting a suitable approach for species delimitation has been difficult, especially in species complexes (Pinheiro et al. 2018). Identifying discontinuities among incipient stages of divergence, which are commonly found in species complexes, demands great effort and a multidisciplinary approach to assess different sources of evidence supporting species limits (Carstens et al. 2013). In this context, estimates based on independent sources of data such as morphology, cytogenetics, anatomy, ecology, and genetics have been used to achieve a better resolution in species delimitation (Domingos et al. 2014; Alvarado-Sizzo et al. 2018; Denham et al. 2019). In particular, phenotypes and geographic distributions can be a starting point for hypotheses about species circumscriptions (Solís-Lemus et al. 2015; Luo et al. 2018).

Species delimitation methods based on multilocus data and the multispecies coalescent model (MSC) compare the probability of species trees with different numbers of operational taxonomic units (OTUs) to identify optimal partitions for the data (Ence and Carstens 2011). Highly fragmented systems impose critical caveats on such methods, potentially resulting in the oversplit of existing diversity, and thus, highly subdivided entities (Sukumaran and Knowles 2017). Jackson et al. (2017) proposed the incorporation of the genealogical divergence index (*gdi*) as an attempt to reduce the delimitation of population structure, instead of species limits. Furthermore, the developers of the Bayesian delimitation method implemented in BPP (for Bayesian Phylogenetics and Phylogeography; Yang and Rannala 2014) described an approach to integrate *gdi* on the outputs of their software and showed that such a strategy was efficient in mitigating the effect of over splitting (Leaché et al. 2018).

However, Rannala and Yang (2020) raised a number of concerns on the use of *gdi*, such as the large interval of *gdi* values (0.2 to 0.7) where delimitation is ambiguous, and misleading results when there are huge differences among the population sizes of the putative species. Model selection methods using simulated datasets under competing delimitation hypotheses are a promising tool for taxa that have potentially experienced a complex evolutionary history, including recurrent gene flow and demographic fluctuations. These methods also allow us to easily test for different assignments and topologies of the analyzed samples into populations/species, a procedure that is not straightforward with other approaches (but see Leaché et al. 2014). Examples of such approaches applied for species delimitation include approximate Bayesian computation (ABC; Camargo et al. 2012) and approximate likelihood analysis (Morales et al. 2017). ABC represents a class of flexible likelihood-free algorithms for performing Bayesian inference (Beaumont et al. 2002). The principle behind the method relies on the massive simulation of genetic data using parameter values drawn from a prior distribution, followed by the calculation of summary statistics (SuSt) for each simulation. SuSt are values generated from the raw genetic data, which are expected to capture important information to differentiate among the simulated models. Simulations that produce genetic variation patterns (SuSt) close to the observed data are retained to form an approximate sample from the posterior distribution.

A number of machine learning methods are increasingly being applied to species delimitation. The applied strategies include coupling ABC with random forest (Smith and Carstens 2020), use a Support Vector Machine on SuSt (Pei et al. 2018) and using different unsupervised machine learning approaches (Derkarabetian et al. 2019). Another promising approach recently developed for demographic model selection, based on deep learning image classification, uses convolutional neural networks (CNNs) to retrieve as much information as possible from genetic datasets by converting them to images without requiring user-specified SuSt (Flagel et al. 2019). Simulations have shown that CNNs maximize the use of available information from the data and increase the ability to distinguish divergent evolutionary histories, frequently overcoming the limitations of other traditional approaches (Flagel et al. 2019). Despite the recent use of CNNs in population genetics (Flagel et al. 2019; Sanchez et al. 2020) and phylogeography (Souza et al. 2019; Oliveira et al. 2020; Fonseca et al. 2021), this approach has not previously been tested in the context of species delimitation using genetic data.

Here, we introduce an approach for species delimitation that combines the use of MSC-based methods with model selection via CNN (detailed in Fig. 1). We compared our model selection method with ABC to evaluate distinct species delimitation hypotheses in the cactus species complex *Pilosocereus aurisetus*, currently arranged in two subspecies. We chose the *P. aurisetus* complex because (1) it presents a controversial taxonomy, and (2) it has a sky-island distribution, which imposes difficulties on MSC-based methods (Leaché et al. 2018). We also validated our method by following this same procedure using a recently published dataset from two *Drosophila* species pairs (Campillo et al. 2020), which have information about pre- and postzygotic reproductive isolation and do not have controversial taxonomy.

**Figure 1.**
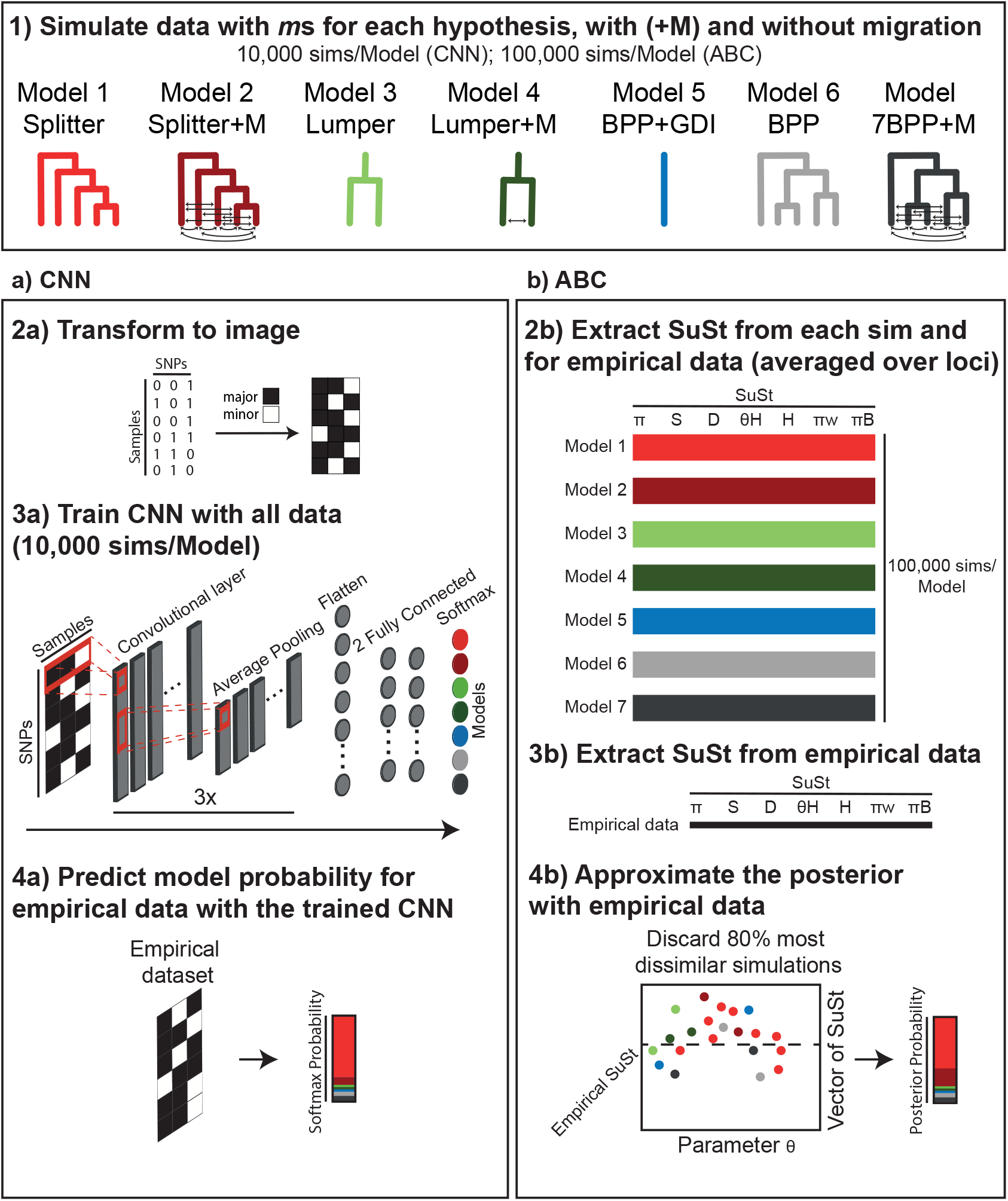
A general protocol for species delimitation in the *P. aurisetus* species complex, comparing the adopted CNN and ABC approaches. The first step for both CNN and ABC was to (1) simulate genetic data for each delimitation hypothesis (10,000 simulations per model in CNN and 100,000 in ABC), with and without gene flow among lineages (+M models). Then, for the a) CNN procedure, we (2a) convert the simulated data to images, with black pixels representing the major and white pixels representing the minor frequency alleles for each segregating site; (3a) use each converted simulation to train a neural network that recognize simulations generated from each model; (4a) predict model probability on our empirical data, using the trained CNN. In our b) ABC approach, we (2b) extract summary statistics (SuSt) averaged over loci for each simulation and (3b) for the empirical data; and (4b) obtain the Posterior Probability of each model by approximating the posterior (retaining only the 20% most similar simulations).

## Material and Methods

### The Study System

The taxon *P. aurisetus* is a puzzling system regarding species delimitation. The taxonomy of *P. aurisetus* complex has been unstable over the years, which is a common attribute in the family Cactaceae, probably due to convergent and parallel evolution causing the absence of clear diagnostic characters for some groups (Hernández-Hernández et al. 2011; Copetti et al. 2017). The morphological variation observed within this taxon has led to repeated taxonomic evaluations, with species and subspecies being recurrently synonymized and re-established (Zappi, 1994; Taylor and Zappi, 2004; Hunt et al., 2006). Another complicating issue is the naturally fragmented sky-island distribution of *P. aurisetus*. The taxon occurence is restricted to the mountaintops of the Espinhaço Mountain Range in eastern Brazil and is a typical element of the *campo rupestre* landscapes (Fig. 2). The *campo rupestre* is a rock outcrop vegetation harboring outstanding species richness and endemism in South American ancient mountaintops (Miola et al. 2021). In line with this sky-island vegetation system, *P. aurisetus* shows high intraspecific genetic structure and restricted or absent gene flow, even among neighbor populations (Bonatelli et al. 2014; Perez et al. 2016a, b). Although the details of the reproductive biology of *P. aurisetus* is unknown, the flower and fruit characteristics are similar to other congeneric taxa, which are predominantly pollinated by bats (Rocha et al. 2019) and seed-dispersed by birds and bats (Vázquez-Castillo et al. 2019). Currently, two subspecies are recognized based on differences in the size and diameter of stems and the color and density of hairs on flower-bearing areoles: the more widespread *P. aurisetus* (Werderm.) Byles & Rowley subsp. *aurisetus*, and *P. aurisetus* subsp. *aurilanatus* (Ritter) Zappi, which is limited to only a few populations restricted to a disjunct mountain (Serra do Cabral) west of the main Espinhaço Mountain Range. We also included plants from three heterotypic synonyms of *P. aurisetus* subsp. *aurisetus* (Fig. S1).

**Figure 2.**
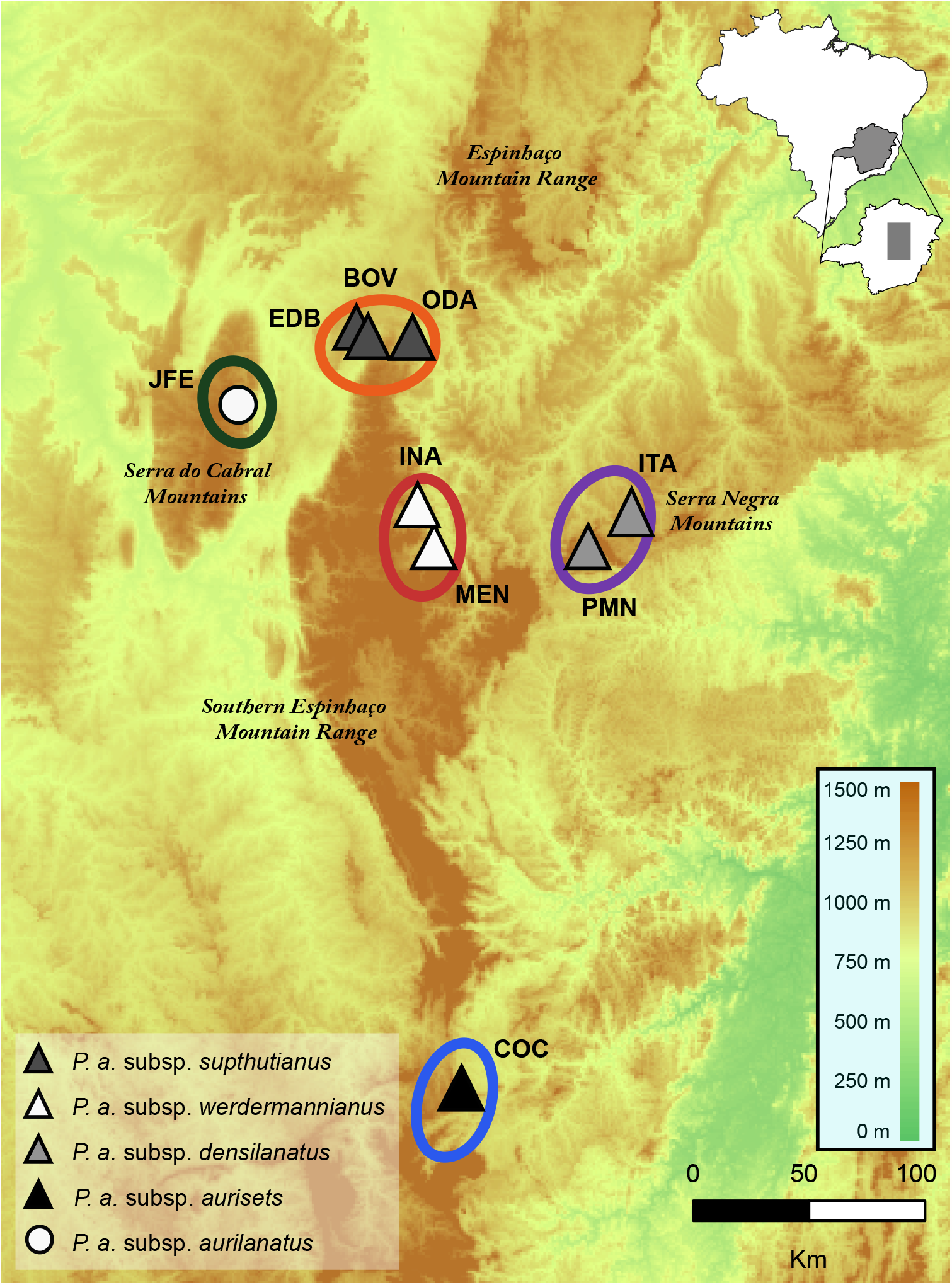
(A) Geographic distribution of the sampled localities and elevational range in eastern Brazil (codes according to Table 1). Symbols (triangles and circle) represent the currently recognized *P. aurisetus* subspecies, and gray shades represent heterotypic synonyms of *P. aurisetus* subsp. *aurisetus* (in parenthesis). The delimited species are shown with lines colored according to the delimited entities in Fig. 3.

The heterotypic synonym *P. aurisetus* subsp. *densilanatus* (Ritter) Braun & Esteves occurs on the easternmost side of the Serra Negra Mountains and is distinguished by their densely wooly stems reaching up to 4 cm in diameter. *P. aurisetus* subsp. *supthutianus* (Braun) Braun & Esteves is a northern outlier of the species that also presents densely hairy stems, which can reach more than 5 cm in diameter. Finally, *P. aurisetus* subsp. *werdermannianus* (Buining & Brederoo) Braun & Esteves occurs in the central range of the *P. aurisetus* distribution and shows more slender stems, fewer ribs and sometimes green epidermis.

**Table 1.** Sampling localities for *P. aurisetus* species complex and their assignment to operational taxonomic units according to three different species delimitation hypotheses (‘splitter’, ‘lumper’, ‘BPP_noGDI’ and ‘BPP_GDI’).

| Taxon/Locality | Code | N | Longitude | Latitude | Delimitation hypothesis |  |  |  |
| --- | --- | --- | --- | --- | --- | --- | --- | --- |
|  |  |  |  |  | ‘splitter’ | ‘lumper’ | ‘BPP_noGDI’ | ‘BPP_GDI’ |
| *P. a. subsp. aurisetus* |  |  |  |  |  |  |  |  |
| Cocais-MG | COC | 4 | -43,450000 | -19,866667 | 1 | 1 | 1 | 1 |
| Itamarandiba-MG (a) | ITA | 4 | -42,935972 | -18,001944 | 2 | 1 | 2 | 1 |
| Pedra Menina-MG (a) | PMN | 4 | -43,044500 | -18,136250 | 2 | 1 | 2 | 1 |
| Olhos D’Água-MG (b) | ODA | 4 | -43,622361 | -17,438833 | 3 | 1 | 3 | 1 |
| Engenheiro Dolabela-MG (b) | EDB | 4 | -43,793667 | -17,451417 | 3 | 1 | 3 | 1 |
| Bocaiúva-MG (b) | BOV | 4 | -43,755833 | -17,545833 | 3 | 1 | 3 | 1 |
| Mendanha-MG (c) | MEN | 4 | -43,536583 | -18,118917 | 4 | 1 | 2 | 1 |
| Inhaí-MG (c) | INA | 4 | -43,605833 | -17,146444 | 4 | 1 | 2 | 1 |
| *P. a. subsp. aurilanatus* |  |  |  |  |  |  |  |  |
| Joaquim Felício-MG | JFE | 4 | -44,178333 | -17,757833 | 5 | 2 | 4 | 1 |

(a) (b) and (c) indicate the samples from the heterotypic synonyms *P. a. subsp. densilanatus*, *P. a. subsp.*
*supthutianus* and *P. a. subsp. werdermannianus*, respectively.

### Sampling and DNA extraction

We sampled four individuals in each of nine localities in eastern Brazil, totaling 36 sampled individuals and covering the entire known distribution of the *P. aurisetus* species complex (Fig. 2; Table 1). Hereafter, we refer to these sampled localities as populations justified by the clear limits of the *P. aurisetus* habitat patches and the marked genetic structure in this taxon (Bonatelli et al. 2014). Root tissue was stored in silica gel and then transferred to a −80°C freezer until DNA extraction. We maintained a distance of approximately 10 m between sampled individuals to avoid collecting clones. Total DNA was extracted using the Extract All kit (Applied Biosystems) and purified with 95% ethanol and 3 M sodium acetate buffer. The DNA concentration was measured with a Qubit fluorometer (Invitrogen).

### Microfluidic PCR and sequencing

Massive parallel target amplification of 41 loci in 36 samples was carried out in microfluidic PCR reactors (Access Array System, Fluidigm). The selected loci included 26 anonymous nuclear markers developed for *P. aurisetus* (Perez et al. 2016c) and another five nuclear, eight plastid and two mitochondrial genic regions selected from the available data in GenBank (Supplementary Table S1). The development of these new markers was based on available sequences from Cactaceae or from other related plant family species to detect conserved regions for primer design. All primers were developed with PRIMER3 v4.0.0 (Untergrasser et al. 2012), with parameters suggested by the Fluidigm Assay Design Group as follows: primer size from 18–23 bases, annealing temperature between 59°C and 61°C, maximum poli-X of 3. The selected markers were synthesized according to Fluidigm specifications and applied to a single 48×48 access array chip with reactions performed according to the manufacturer’s protocol. The obtained PCR products were pooled and purified with 0.6X AMPure beads (Agencourt). The quality and amplification range were visualized in a BioAnalyzer, and the concentration was estimated via real-time PCR using KAPA qPCR (Kapa Biosystems). The samples were subjected to a MiSeq (Illumina) 300 bp paired-end run along with 156 other samples from other projects.

After sequencing, a first filtering step was performed with cutadapt (Martin 2011), excluding sequences smaller than 60 bp and removing 3’ bases with a PHRED score <Q22. PHRED quality for all reads was then visualized in FastQC (http://www.bioinformatics.bbsrc.ac.uk/projects/fastqc) and graphically compiled in MultiQC (http://multiqc.info/). Reference sequences for each marker were generated from the alignments used to design the primers. Mapping reads for each marker was performed with BWA-MEM (Li 2013), which is suited for paired-end long reads with indels, while SAMtools (Li et al. 2009) was used to retain only sequences that were mapped with high fidelity. Single-nucleotide polymorphism (SNP) calling was carried out with GATK (McKenna et al. 2010) by transforming low-quality bases to missing data, identifying possible indels with realignment, applying polymorphism filters for quality and coverage, and generating FASTA files for each sample. To confirm the obtained polymorphisms, each detected SNP was then mapped against reference files for each marker that were built from reads from each analyzed sample. The phasing of SNPs in nuclear markers was achieved with a Python script, modified from Harvey et al. (2016). The aligned sequences from each marker were obtained with MAFFT v.7 (Katoh and Standley 2013) with default parameters.

Recombination within each marker was tested with the DSS method (McGuire and Wright 2000) implemented in TOPALi v2 (Milne et al. 2008) and topological incongruences were tested in KDETrees (Weyenberg et al. 2014). The most likely substitution model for each locus was obtained with PartitionFinder v.1.1.1 (Lanfear et al. 2012).

### Species Tree and Coalescent Delimitation

The phylogenetic relationships among populations were recovered with a species tree in StarBEAST v2.1.3 (Bouckaert et al. 2014), assigning the sequences from each population to a single OTU. For all markers, we used the selected substitution model (Table S1) and a Relaxed clock Log-Normal, with the mean sampled from a broad lognormal distribution (according to Perez et al. 2016b). We performed three runs with 5 x 10^8^ MCMC iterations each, sampling trees every 10^4^ generations with 75% burn-in. Convergence was observed in TRACER 1.6 (Rambaut et al. 2014) and a Maximum Clade Credibility (MCC) tree with median heights was obtained in TreeAnnotator (Drummond and Rambaut 2007).

We used the obtained species tree topology and the genomic sequences obtained from microfluidic PCR as input to test the species limits in *P. aurisetus* complex with the method (A10 analysis) implemented in BPP v4.2.9 (Yang and Rannala 2014; Flouri et al. 2018). For this analysis, we concatenated all mitochondrial (*cox1* and *cox2*) and chloroplast (*atpB-rbcL, petL-psbE* and *rpl16*) markers. The BPP method uses a reversible jump Bayesian (rjMCMC) framework to estimate the posterior probability of a species split at each node of the phylogeny. To ensure convergence, five independent runs were performed to estimate the best delimitation scheme, with parameters selected according to the values suggested by Leaché et al. (2018) combined with previous estimates for the species from Perez et al. (2016a). *θ* and τ priors followed an inverse gamma (IG) distribution with ?=3 and a β of 0.01 and 0.002, respectively (hereafter BPP ‘specific prior values’). We evaluated the potential effect of priors in the delimitation results by also performing the same number of runs with broad prior values *θ* ∼ IG(3, 0.002) and τ ∼ IG(3, 0.03) as suggested in the BPP manual (hereafter ‘diffuse prior values’). We considered a node as valid only if both analyses support its existence with a posterior probability (PP) of 0.95 or higher. The runs were carried out for 10^5^ iterations sampled every five steps after discarding the first 10^4^ iterations as burn-in. The obtained delimitations were mapped to the tips of the species tree topology. To evaluate the potential effect of tree uncertainty in species delimitation, we also performed additional BPP runs using the same strategies described above, but with a joint species delimitation and species tree inference (A11 analysis). Also, as a more conservative alternative, we evaluated our BPP results using the *gdi* index, as suggested by Leaché et al. (2018). We considered the threshold values suggested by Jackson et al. (2017) and Leaché et al. (2018), in which pairwise *gdi* values smaller than 0.2 indicate a single species and above 0.7, two species.

### Comparing Delimitation Hypotheses with Deep Learning and ABC

We adopted an integrative approach to establish putative species limits by considering information on the *P. aurisetus* complex from different sources and analyses, as suggested by Carstens et al. (2013). For that, we compared the delimitation hypotheses generated by the BPP results (with and without the *gdi* index), as well as two morphological hypotheses based on one lumper and one splitter taxonomic arrangement (Table 1; Fig. 1). The genetic hypotheses, based on the BPP results, were composed of one species when *gdi* was considered and four species when it was not incorporated (see results below); the ‘splitter’ morphological hypothesis, considering both currently accepted and previously synonymized taxa, was composed of four species; and the ‘lumper’ morphological hypothesis, considering only the currently recognized subspecies, was composed of two species (Hunt et al. 2006). To test these competing delimitation hypotheses, we considered scenarios including the number of species and the potential migration between the delimited entities. Therefore, seven scenarios (Fig. 1) were tested: (1) ‘splitter’, (2) ‘splitter’ combined with migration, (3) ‘lumper’, (4) ‘lumper’ with migration, (5) BPP combined with *gdi* (BPP_GDI), (6) BPP without *gdi* (BPP_noGDI), and (7) BPP_noGDI with migration. As the BPP_GDI scenario consists of a single species, an additional model including migration was not necessary. We performed model comparison using both a deep learning approach based on CNN and a regular ABC method. For these two strategies, we simulated genetic datasets under each scenario with a modified version of the scripts from Perez et al. (2016a). To simplify our simulations and avoid using an overly heterogeneous dataset, we included only nuclear markers and simulated data mirroring sample sizes with the same number of segregating sites per loci of our empirical dataset. For the deep learning approach, 10,000 simulated datasets per model were converted to images that were used to train a CNN to recognize simulations generated by different scenarios (for details on the simulation and training procedure, see Supplementary File 1). The comparison of delimitation scenarios using regular ABC analysis (Fig. 1) was performed following the approach of Perez et al. (2016a) with 100,000 simulations per model conducted under the same conditions used for the CNN (details in Supplementary File 1).

To validate our newly proposed deep learning method for species delimitation, we also carried out the same analyses using a dataset recently published by Campillo et al. (2020). These authors compared the results of reproductive isolation and coalescent-based (BPP) delimitation methods to infer species boundaries in several species pairs of the genus *Drosophila*, a classic model system in speciation research. We focused our approach on data from two species pairs of the Melanogaster group: *D. melanogaster* - *D. sechellia* and *D. melanogaster* - *D. simulans*. We chose these species because they are widely accepted and corroborated by both BPP and reproductive isolation metrics (Campillo et al. 2020). Furthermore, while the *D. melanogaster* - *D. sechellia* pair is allopatric, the *D. melanogaster* - *D. simulans* pair is sympatric, allowing us to explore whether geographic context might affect our ability to identify these species. Therefore, we tested a model with all conspecifics against a model in which the specimens were divided into two species, which was the expected outcome according to the currently supported classification. As with *P. aurisetus*, our simulations were based on empirical sample sizes and the number of segregating sites (details in Supplementary File 1), with priors based on the values suggested by Campillo et al. (2020).

## Results

### Microfluidic PCR and sequencing

Four markers showed no amplification in any sample (*PaANL_*46, *petA, rpS16*, and *ycf1*). To further minimize missing data and ensure that at least one individual was sequenced from each population, we discarded 16 loci (*PaANL_*8, *PaANL_*10, *PaANL_*50, *PaANL_*82, *PaANL_*123, *PaANL_*134, *PaANL_*142, *PaANL_*155, *PaANL_*160, *PaANL_*187, *PaANL_*196, *its2, ppc, nhx1, psbA-psbB, psbC*). No recombination was detected at any of the remaining loci, but kdetrees result suggested an incongruent topology in the *isi1* marker, which was removed. Therefore, the final dataset included 20 markers that were considered in the subsequent analyses for *P. aurisetus* (*PaANL_*15, *PaANL_*17, *PaANL_*28, *PaANL_*35, *PaANL_*80, *PaANL_*87, *PaANL_*96, *PaANL_*126, *PaANL_*140, *PaANL_*147, *PaANL_*164, *PaANL_*165, *PaANL_*182, *PaANL_*205, *apk1, atpB-rbcL, petL-psbE, rpl16, cox1* and *cox2*), with a total of 8,908 bp.

### Species Tree and Coalescent Delimitation

The StarBEAST species tree recovered most nodes with high support, especially the ones defining previously and currently valid subspecies (Fig. 3). Population JFE of *P. aurisetus* subsp. *aurilanatus* was the most external clade. Other populations were arranged in two subclades. One of them contained populations in the center (INA and MEN, previously classified as *P. a*. subsp. *werdermannianus*; ITA and PMN, formerly *P. a*. subsp. *densilanatus*) and south (COC) of the *P. a*. subsp. *aurisetus* distribution. The second subclade contained populations from northern distribution (EDB, BOV and ODA, formerly classified as *P. a*. subsp. *supthutianus*).

**Figure 3.**
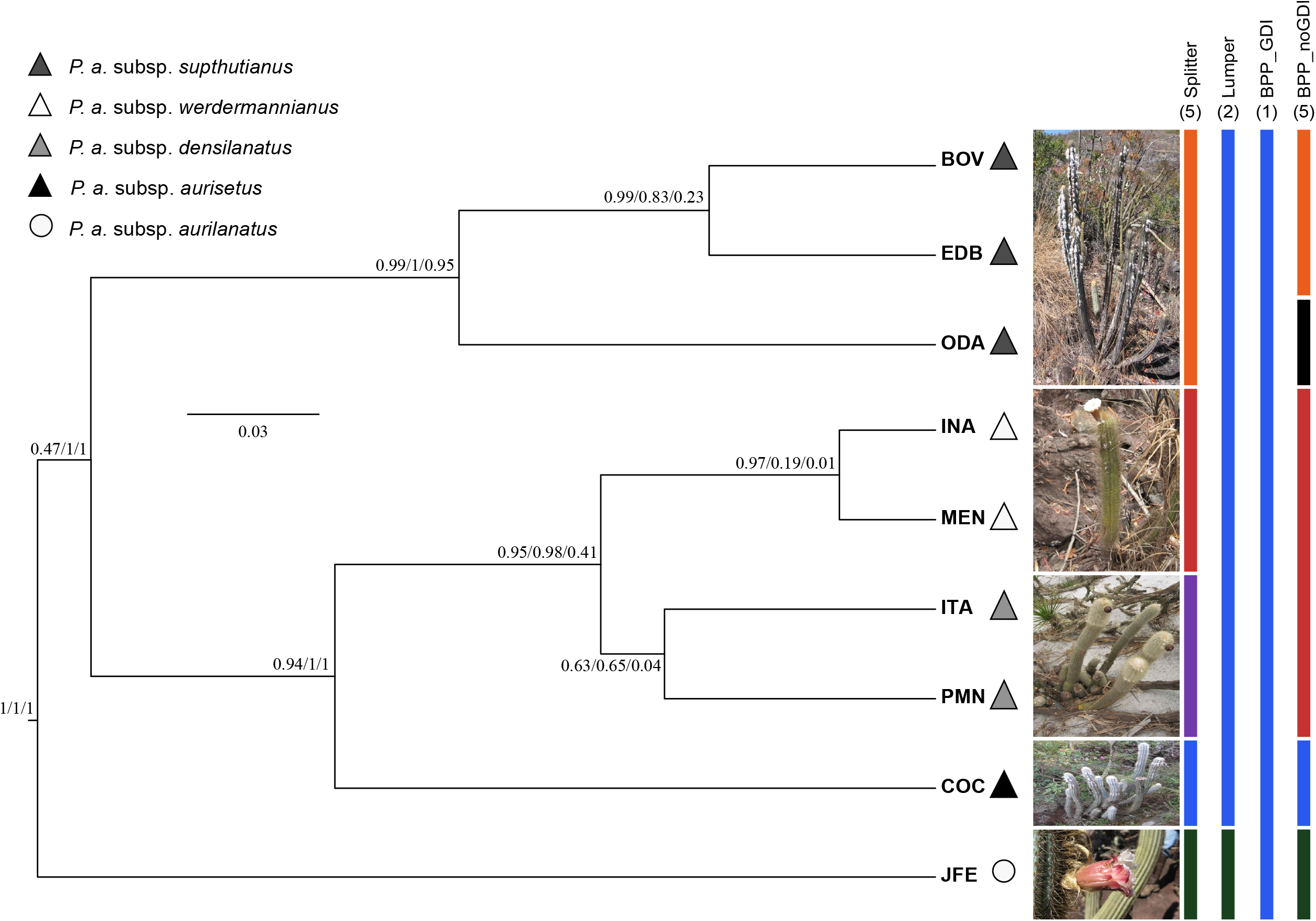
Species delimitation results are represented as bars at the tips of the topology obtained in BPP. Values in parentheses above each bar present the number of species for each hypothesis. The numbers at the nodes represent Beast PP/BPP ‘specific’/and ‘diffuse’ prior values. Symbols (triangles and circles) represent both currently recognized and synonymized (in parenthesis) *P. aurisetus* taxa. A representative of each inferred species is shown, next to the limits according to the ‘splitter’ hypothesis.

All BPP runs (A10 analysis) under each prior set showed the same results, therefore we provided the mean posterior probability of each node (Fig. 3). Results with ‘specific’ and ‘diffuse’ priors were similar, though the latter supported a lower number of species. The congruent delimited species were *P. a*. subsp. *aurisetus* (COC), *P. a*. subsp. *supthutianus* populations EDB and BOV separated from ODA and the currently recognized subspecies *P. aurisetus* subsp. *aurilanatus* (JFE). The only exception was for the node separating *P. a*. subsp. *werdermannianus* (INA and MEN) and *P. a*. subsp. *densilanatus* (ITA and PMN), which was validated (PP=0.98) only in the ‘specific’ but not (PP=0.41) in the ‘diffuse’ prior set. To be conservative, we decided to collapse this node, considering them as a single unit. Therefore, BPP resulted in five delimited species that coincided with the ‘splitter’ hypothesis, except for the OTUs composed by populations once recognized as *P. a*. subsp. *werdermannianus* and *P. a*. subsp. *densilanatus*, which were combined, and for population ODA that was separated from the other *P. a*. subsp. *supthutianus* populations (BOV and EDB) (Fig. 3). BPP results from the joint species delimitation and species tree (A11 analysis) were similar, with the ‘specific’ prior runs resulting in more delimited units (7 - assigning each population to a single species, expect for populations INA, ITA and MEN that were collapsed), while the runs with the ‘diffuse’ priors recovered the same five species described above (BPP_noGDI in Fig. 3). When we analyzed our BPP results with the *gdi* index, all estimates were below 0.7 and only three (*P. aurisetus* subsp. *aurilanatus* - JFE; *P. a*. subsp. *aurisetus* - COC; and *P. a*. subsp. *supthutianus* - ODA, BOV and EDB) had values between 0.2 and 0.7 (Fig. S2). Therefore, to be conservative, we decided to collapse all populations in a single species for this hypothesis.

### Delimitation Hypotheses Comparison

After 250 epochs, our CNN showed accuracies of 96.81% and 92.49% for the training and validation sets, respectively. Some degree of overfitting was observed when we plotted the accuracy of the training and validation sets throughout the epochs. However, this effect decreased inversely proportional to the number of evaluated simulations (Fig. S3). Our cross-validation procedure, using a test set of simulations not evaluated during training, also showed that increasing the number of simulations resulted in accuracy improvement for all models (Fig. 4). It was also possible to observe that CNN presented a higher proportion of simulations that were correctly predicted in relation to their model than the ABC results, with all models being correctly assigned with more than 80% accuracy when 10,000 simulations per model were used (Fig. 4 and S4). The percentages of correct predictions with the ABC approach were all higher than 70% when 100,000 and 10,000 simulations per model were used, except for the models ‘splitter’ and ‘BPP_noGDI’ that showed ∼50% accuracy (Fig. 4 and Fig. S4). These two latter models also showed the highest levels of confusion with each other (Fig. S4). The ‘splitter’ hypothesis was selected with a softmax probability of 99.9% (Table 2) when the empirical data were submitted to the trained CNN, and this result was consistent even when the order of SNPs was shuffled (Supplementary File 1). The ABC approach showed similar results, also selecting the ‘splitter’ hypothesis (PP=0.962). Further, when we plotted the simulated and empirical genotypes and the ABC posterior with a PCA (Fig. S5) the empirical data were located inside the cloud of simulations from the ‘splitter’ model.

**Figure 4.**
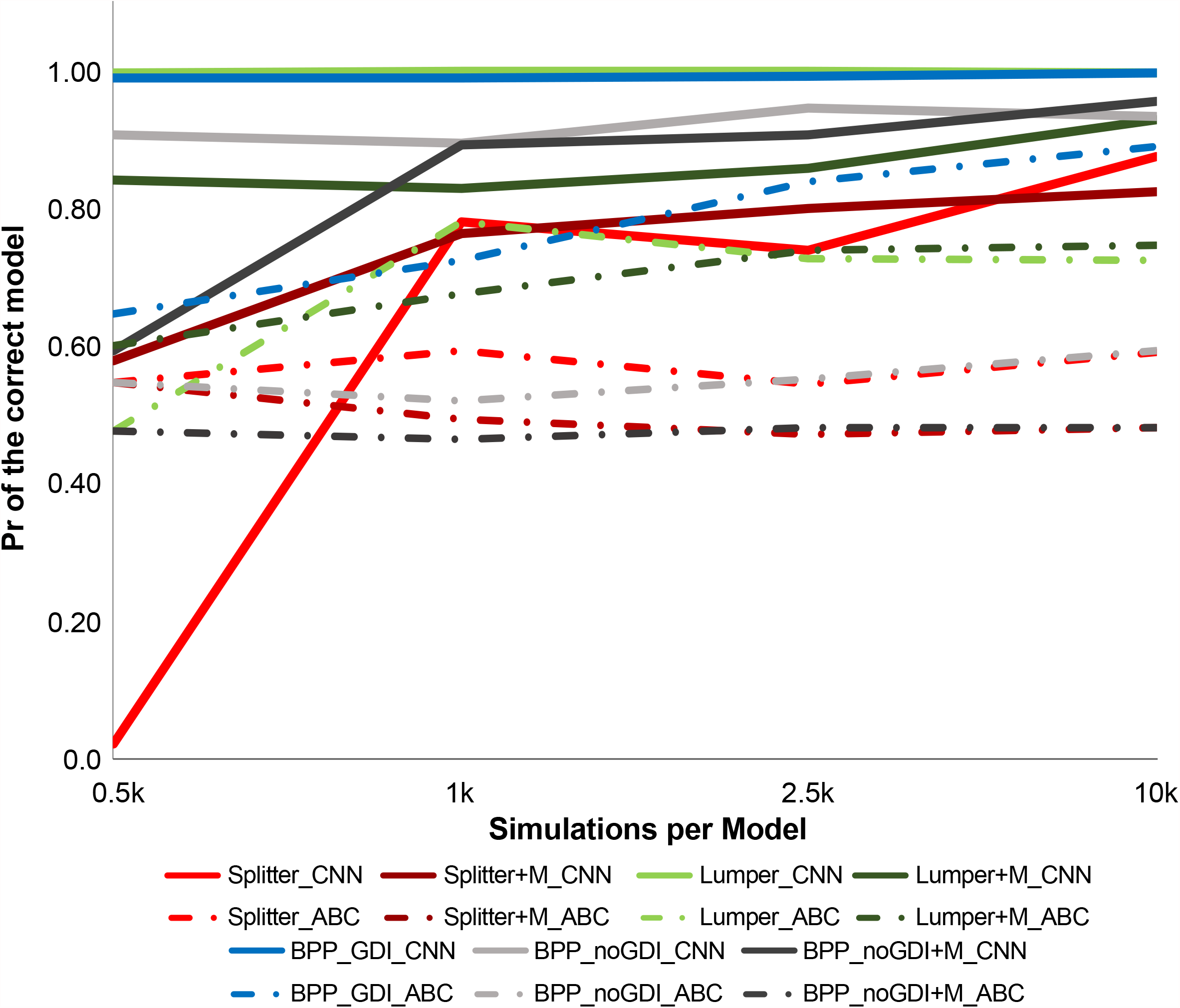
Cross-validation test to compare the performance of different model comparison methods (CNN - filled lines and ABC - dashed lines) and numbers of simulations. Each point represents the probability of choosing the “correct” model (i.e., the model from which the data were simulated) over incorrect models. Each color represents a different scenario, including (+M) or not migration between the OTUs.

**Table 2.** Model selection results for the *P. aurisetus* species complex and the two *Drosophila* species pairs using empirical data. CNN values are softmax probabilities for each scenario, while posterior probabilities are shown for ABC. The scenario with the highest probability for each method is shown in bold.

| Scenario | CNN | ABC |
| --- | --- | --- |
| *P. aurisetus* |  |  |
| Splitter | **0.999** | **0.962** |
| Splitter+Migration | 0.000 | 0.002 |
| Lumper | 0.000 | 0.000 |
| Lumper+Migration | 0.000 | 0.000 |
| BPP+GDI | 0.000 | 0.000 |
| BPP_noGDI | 0.001 | 0.033 |
| BPP_noGDI+Migration | 0.000 | 0.003 |
| *D. melanogaster_D.simulans* |  |  |
| One Species | 0.008 | 0.064 |
| Two Species | **0.992** | **0.936** |
| *D. melanogaster_D. sechellia* |  |  |
| One Species | 0.051 | 0.474 |
| Two Species | **0.949** | **0.526** |

The *Drosophila* dataset also pointed to a higher accuracy of CNN over ABC. The cross-validation tests indicate a correct assignment of CNN simulations just above 78% and 81% in the *D. melanogaster-D simulans* and *D. melanogaster-D. sechellia* pairs, respectively (Fig. S4). The accuracy of ABC was lower, reaching values slightly higher than 72% and 68% for the same species pairs (Fig. S4). When the empirical data were used, both methods supported the hypothesis of two species for the *D. melanogaster-D simulans* pair, showing a probability of 99.2% in CNN and 93.6% for ABC. In *D. melanogaster-D. sechellia*, both methods also pointed to two species with a probability of 94.9% for CNN and 52.6% for ABC (Table 2).

## Discussion

### Deep learning in coalescent-based species delimitation

The advantages of the simulation approaches adopted here (CNN and ABC) over explicit statistical methods are their ability to test models with complex demographic scenarios, such as migration events or size fluctuations. In addition, these approaches enable us to easily test for different assignments of samples into OTUs (but see Leaché et al. 2014) and even compare hypotheses relying on distinct topologies. Here, we compared hypotheses derived from two previous taxonomic arrangements with the results we obtained from BPP. For each hypothesis, we also evaluated whether gene flow among the delimited entities was present, totaling seven evaluated scenarios (Fig. 1). We decided not to include even more complex scenarios (e.g. by incorporating demographic fluctuations), as evaluating a high number of scenarios can negatively impact the performance of the methods applied, especially in ABC (Pelletier & Carstens 2014). Indeed, our comparison of the two approximate approaches suggests that deep learning (CNN) showed a higher capacity to distinguish among the simulated demographic scenarios, outperforming ABC (Fig. 4 and S4). It is important to highlight that CNN achieved this performance with ten times less simulations and much smaller running times (∼22 minutes to train the network in a Nvidia K80 GPU with an Intel Xeon 2.30GHz CPU) than ABC (∼17 days to perform cross-validation and 19 minutes for model selection in an Intel Xeon 2.40GHz CPU; see Table 3 for a full comparison of running times for each step). The results from the *Drosophila* dataset also supported a higher accuracy of CNN compared to ABC (Fig. S4). Moreover, when the empirical data was evaluated, CNN recovered two species for both Drosophila species pairs with higher probability, especially for *D. melanogaster-D. sechellia* (Table 2). These results validated our CNN approach, as they are in agreement with our expectations based on Campillo et al. (2020).

**Table 3.**
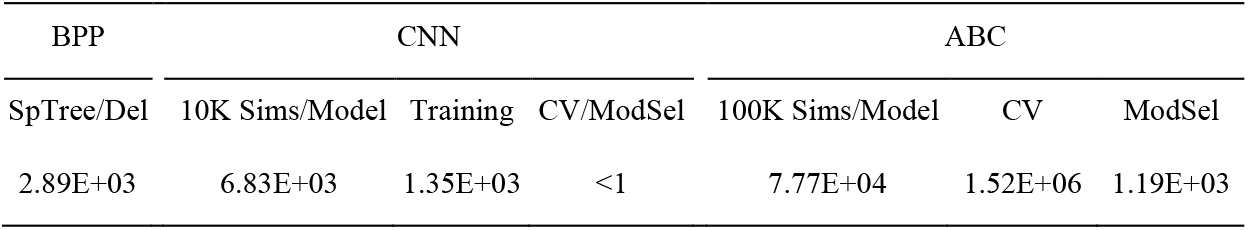
Computational time to estimate the species tree and delimitation (SpTree/Del) in BPP, run 10,000 (10K) simulations per model (Sims/Model) for CNN and 100,000 (100K) for ABC, train the neural network (for CNN), and perform cross validation (CV) and model selection (ModSel) for CNN and ABC approaches, in seconds.

Regarding the *P. aurisetus* dataset, both CNN and ABC approaches selected the ‘splitter’ model without migration as the most likely scenario with high PP (Table 2). The main difference between CNN and ABC functioning is the ability of the former to take information directly from SNP matrices without using SuSt, which is required by the latter. Moreover, while ABC relies on a rejection step that discards most of the simulations and leaves only a small proportion that are more similar to the empirical data (Csilléry et al. 2012), the CNN uses information from the whole set of simulations to learn how to distinguish the concurrent scenarios (Schrider and Kern 2018). The inferior performance of the ABC analyses could also be related to the choice of SuSt with overly high dimensionality (Robert et al. 2011; but see Kousathanas et al. 2016) or with little information to distinguish among the tested models. To reduce dimensionality, we adopted the suggestion of Perez et al. (2016a) applying a PCA transformation of the raw SuSt and taking only the most explanatory axes necessary to ensure at least 99% of the variance in the data. Other strategies usually applied to reduce dimensionality (reviewed in Prangle 2015) include partial least squares regression (Wegmann et al. 2009), linear regression (Fearnhead and Prangle 2012) or boosting (Aeschbacher et al. 2012).

Although our CNN results were very encouraging, we note that care must be taken in interpretation, as deep learning techniques can suffer from overfitting (Nguyen et al. 2015; Ponti et al. 2017). To evaluate overfitting, we observed the history plot of the accuracy throughout the epochs (Fig. S3) and adopted a cross-validation approach based on a test set, which consisted of simulated data that were not evaluated during training. The results pointed to high accuracy (Fig. 4 and Fig. S4), which is in agreement with the similarity of our empirical and simulated data (Fig. S5). It is important to note that our CNN approach is very flexible and can potentially be combined with other recently developed machine learning applications. For instance, our CNN approach can be used to compare models and estimate parameters by using the CNN predictions (Mondal et al. 2019) or combining them with SuSt (Sanchez et al. 2020) to perform ABC. Some machine learning approaches have also shown promising results for the specific field of species delimitation, and they could also be combined or compared to our CNN method (Pei et al. 2018; Derkarabetian et al. 2019; Smith and Carstens 2020).

### Delimiting Species in <u>P. aurisetus</u>

*Pilosocereus aurisetus* populations diverged very recently in the last one million years (Perez et al. 2016a). Delimiting species with recent diversification has been considered challenging, and the main difficulty arises from the short time for species to accumulate diagnostic characteristics that allow taxonomic distinction (Salicini et al. 2011). A similar effect is observed in genetic data and is commonly referred to as incomplete lineage sorting (ILS). ILS appears when ancestral genetic variants persist as polymorphisms across successive diversification events. This phenomenon has been indicated as a major cause of discordance between gene trees and species trees in Cactaceae (Copetti et al. 2017).

Another issue that imposes additional challenges for species delimitation in *P. aurisetus* complex is its sky-island distribution, constraining gene flow and leading to high population divergence (Bonatelli et al. 2014). Topography-driven isolation in species of the Brazilian *campo rupestre* environments, such as *P. aurisetus*, has been considered an important driver of speciation and high microendemism in these landscapes (Vasconcelos et al. 2020; Zappi et al. 2017). The major concern in this kind of system is that MSC based methods may fail to discriminate between genetic structure associated with population isolation and species boundaries (Jackson et al. 2017; Sukumaran and Knowles 2017; Leaché et al. 2018). Although *gdi* appears as a possible solution for the overspliting tendency of MSC methods (Jackson et al. 2017), mainly in highly structured systems, it is still not clear if it could lead to an opposite effect - over lumping differentiated species. Incorporating *a priori* knowledge on the structure associated with population-level processes likely could overcome this overspliting tendency of MSC methods. An ideal scenario would incorporate data from a plethora of classes, such as morphological, ecological, and genetic data, to recover more robust species boundaries (Pinheiro et al. 2018). Since no single method can discriminate between these processes, the results from species delimitation using multispecies coalescent models (MSC) in sky-island systems should be treated as species hypotheses that need further confirmation from multiple sources (Sukumaran and Knowles 2017).

### *Taxonomic Implications for <u>P. aurisetus</u>* complex

Historically, Cactaceae taxonomy reflected the amateur views, always looking for differences with the objective to describe all possible variation found within and between populations as new species. Zappi (1994), Taylor and Zappi (2004) and Hunt et al. (2006) sought similarities and relationships between species, leading to a lumping approach. The present species delimitation investigation within the *P. aurisetus* complex is discordant from the currently recognized taxonomic diversity of this taxon (Hunt et al. 2006), recovering the ‘splitter’ hypothesis with higher probability than different models. Convergent evolution as well as the level of plasticity commonly found in cactus species (Guerrero et al. 2018) might be related to the disagreement in the number of species delimited by current taxonomy and the detected genetic lineages, as observed within the genus *Pilosocereus* (Bonatelli et al. 2014; Perez et al. 2016b). Furthermore, the ‘splitter’ hypothesis is in line with the sky-island distribution and the pronounced population geographic isolation in this taxon, which agrees with the high incidence of microendemic plant species in the *campo rupestre* landscapes (Miola et al. 2021). Therefore, our results show the need for a taxonomic revision of the *P. aurisetus* complex and provide recommendations for a taxonomic treatment that better reflects the diversity of species. In this context, our ‘splitter’ hypothesis may be used as a guide to investigate new diagnostic characters among the proposed species, contributing to fill gaps in the traditional taxonomy of cactus. Previous genetic studies have identified well-differentiated lineages within *P. aurisetus* complex and related species, with monophyletic groups more frequently being associated with the geographic distribution than the proposed taxonomic boundaries (Fig. 2; Bonatelli et al. 2014; Perez et al. 2016b; Calvente et al. 2016; Khan et al. 2018; Lavor et al. 2019). Different taxonomic circumscriptions have been considered in the past for some of the recovered lineages within the *P. aurisetus* complex that agree with our results. For instance, the populations occurring in the Serra Negra Mountains (ITA and PMN localities), in the center (MEN and INA) and in the northern distribution of the species (EDB, BOV and ODA localities) were previously reported as *P. aurisetus* subsp. *densilanatus, P. aurisetus* subsp. *werdermannianus*, and *P. aurisetus* subsp. *Supthutianus* (Braun and Esteves Pereira 1995), respectively. These subspecies propositions were based on some level of morphological variation that initially seemed rather distinctive, such as the densely wooly stems observed in the ITA populations (Fig. S1). However, Taylor and Zappi (2004) argued that such differences were not sufficiently important to merit recognition as an additional subspecies. These authors reasoned that if this level of morphological difference was to be recognized, the logical extension would be to give similar status to other populations that show slightly divergent characteristics in terms of size, white-wooly stems, and glaucousness.

### Concluding Remarks

Although the splits within species could be an artifact of the MSC, we might recognize previous taxonomic issues that agree with the inferred species in this investigation. *Pilosocereus aurisetus* complex exhibits many biological characteristics that make species delimitation challenging, such as recent divergence, a sky-island distribution and unstable taxonomic history. Here, we sought to investigate the species limits in this complex system and, for the first time, incorporated a deep learning approach to select the best species hypothesis according to our data and simulations. Finally, we stress that all the species hypotheses reported here should be considered, as any of the taxonomic species in fact are hypotheses prone to validation with other methods and sources of information.

## Acknowledgments

This work was supported by grants from the São Paulo Research Foundation (FAPESP) (2015/06160-5 to EMM, 2012/22943-1 to MFP, and 2012/22857-8 to IASB); the National Council for Scientific and Technological Development (CNPq) (03940/2019-0 to EMM, 305301/2018-7 to DCZ); and the Coordination for the Improvement of Higher Education Personnel (CAPES) (Finance Code 001 to MRB). We thank Heidi Utsunomiya and Juliana de Fátima Martinez for laboratory assistance and Gerardus Olsthoorn and Marlon Machado for sampling assistance.

## Data Accessibility

All sequences were deposited in GenBank with accession numbers MZ509667 - MZ510910. All scripts and datasets used are available in GitHub: https://github.com/manolofperez/CNN_spDelimitation_Piloso.

## Author Contributions

M.F.P., I.A.S.B, F.F.F., D.C.Z., N.P.T., and E.M.M. conceived and designed the study. M.F.P. performed the analysis, with critical inputs from I.A.S.B, E.M.M, and F.F.F. M.F.P., I.A.S.B. and M.R.B. drafted the manuscript. D.C.Z. and N.P.T. guided the sampling of the taxonomic complexity of the study group, identified the species, and helped with the fieldwork. All authors participated in discussions and contributed critically to data interpretation and the final version of the text.

