## Supplementary material for "Coalescent-based species delimitation meets deep learning: Insights from a highly fragmented cactus system": Supp File 1

SUPPLEMENTARY MATERIALS

MANOLO F. PEREZ[1,2,¥], ISABEL A. S. BONATELLI[1,3,,¥], MONIQUE ROMEIRO-BRITO[1], FERNANDO F. FRANCO[1], NIGEL P. TAYLOR[4], DANIELA C. ZAPPI[5], EVANDRO M. MORAES[1,*]

*[1]Departamento de Biologia, Universidade Federal de São Carlos, Rodovia João Leme dos Santos km 110, Sorocaba, SP, 18052780, Brazil;*
*[2]Departamento de Genética e Evolução, Universidade Federal de São Carlos, Rodovia Washington Luiz km 235, C.P. 676, São Carlos, SP, 13565905, Brazil;*
*[3]Departamento de Ecologia e Biologia Evolutiva, Universidade Federal de São Paulo, Diadema, SP 09972-270, Brazil;*
*[4]University of Gibraltar, Gibraltar Botanic Gardens Campus, The Alameda, PO Box 843, Gibraltar;*
*[5]Programa de Pós Graduação em Botânica, Instituto de Ciências Biológicas, Universidade de Brasília, PO Box 04457, Brasília, DF, 70910970, Brazil.*
*[¥] Manolo F. Perez and Isabel A. S. Bonatelli contributed equally to this article.*

### Supplementary Text

#### *Simulation details*

We simulated genetic datasets for *P. aurisetus* with a modified version of the scripts from Perez et al. (2016) under each delimitation scenario (Fig. 1 in main text): (1) ‘splitter’, (2) ‘splitter’ combined with migration, (3) ‘lumper’, (4) ‘lumper’ with migration, and (5) BPP combined with *gdi* ‘BPP_GDIgenetic’, (6) BPP without *gdi* ‘BPP_noGDI’, and (7) ‘BPP_noGDI’ with migration. Simulations were performed using the coalescent simulator *ms* (Hudson 2002), using empirical sample sizes and simulating each loci separately, but linking the segregating sites of each loci (fixed by using the -s option), according to the converted empirical data. To convert the empirical data to the same format (*ms*) as the simulations, we used fastaconvrtr (available at https://github.com/CRAGENOMICA/fastaconvtr/). This procedure was performed independently for each loci, and the resulting files were combined after conversion. As this conversion discards missing data, we removed one sample from each of the ITA, PMN, EDB and ODA populations before converting, to retain a higher number of loci. It is important to note that this conversion removes non-binary segregating sites (the final data matrix resulted in of 195 segregating sites). A uniform distribution between 1 and 15 was used for $\theta$, and divergence times for the root were sampled from a uniform distribution between 0.5 and 5 coalescent units. For the models including migration, a uniform prior from 0 to 5 migrants per generation was used. These values were selected according to results from a previous study in *P. aurisetus* (Perez et al. 2016). Subsequent splits in the simulated trees were sampled from uniform priors from the present to the time of the previous split (for details, see scripts provided at GitHub: https://github.com/manolofperez/CNN_spDelimitation_Piloso). We transformed our simulated data into NumPy arrays (images) including specimens as lines and loci as columns. As we do not have information on the polarity of our segregating sites, we coded the genotypes as 0 (black) for the major state and 1 (white) for the minor state. Then, for each simulation, the matrix was transposed, resulting in the allocation of each specimen to a column, while each segregating site occupied a line (Supplementary Text Figure 1).

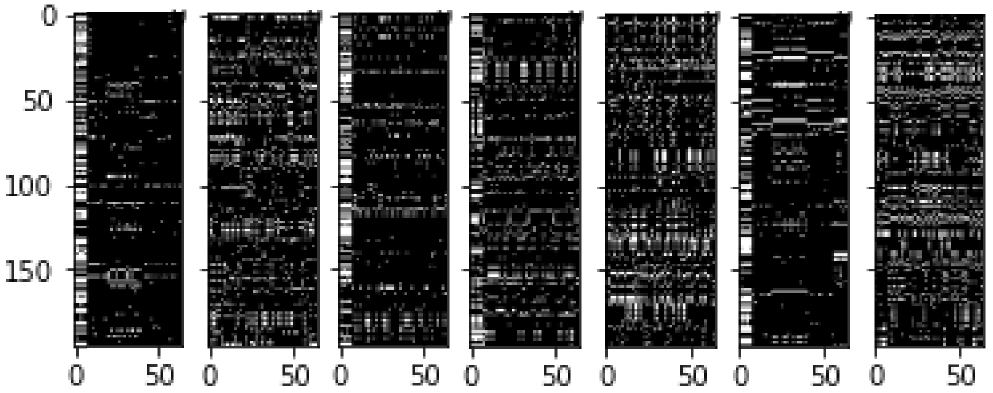


SUPPLEMENTARY TEXT FIGURE 1. Simulated datasets under the seven delimitation scenarios tested. From left to right: (1) ‘splitter’, (2) ‘splitter’ combined with migration,

(3) ‘lumper’, (4) ‘lumper’ with migration, and (5) BPP combined with *gdi* ‘BPP_GDIgenetic’, (6) BPP without *gdi* ‘BPP_noGDI’, and (7) ‘BPP_noGDI’ with migration.

*Deep Learning model selection approach*

The order of the resulting NumPy arrays was then shuffled, and 25% random simulations were separated to be used as the validation set, while the remaining 75% were used as the training data. The resulting dataset was submitted to a convolutional neural network (CNN) based on the architecture suggested by Flagel et al. (2019), with slight modifications (see Fig. 1 in the main text). Three 1D-convolutional layers with a kernel size of 2 (with the first layer containing 250 and the other two, 125 neurons each) were intercalated with average pooling steps. The convolutional layers extracted features of the input image, such as edges, by applying filters (kernels) that slide (convolves) along the input image. The average pooling layers then reduced the dimensionality by recovering the average value of the area covered by the kernel, which extracts dominant features of the data. Thereafter, two fully connected layers with 125 neurons were used to connect all previous neurons and summarize all learned information. Finally, an output layer with neurons corresponding to the scenarios used to simulate the data was employed to compute the probability for each model using a softmax function. Runs were carried out with a minibatch size of 250 for 250 epochs, using rectified linear unit activation functions (i.e., ReLUs; Nair and Hinton 2010) and dropout regularization to avoid overfitting (removing 25% and 50% of neurons at random in each training iteration from the convolutional and densely connected layers, respectively). We used the Adam optimization procedure to update the network weights (Kingma and Ba 2014) and evaluated the network performance with a categorical cross-entropy loss function. To further avoid overfitting, we also used a learning rate reduction when accuracy not improved for more than 20 epochs, together with a checkpoint strategy to retain only the model with the highest accuracy on the validation set. We evaluated the actual accuracy of our trained networks to recognize the scenario from each simulation using a test set of 1,000 simulations per model. This set of independent simulations, not evaluated by the network during the training, allows us to estimate the diagnostic capacity of the network by building a confusion matrix of the recovered model for each analyzed simulation. After that, the trained network was used to predict the most likely scenario using the empirical data as input (Supplementary Text Figure 2). We also tested the impact of shuffling the order of segregating sites, which returned similar results (see the results in the Jupiter Notebook "CNN_Piloso.ipynb" at: https://github.com/manolofperez/CNN_spDelimitation_Piloso).

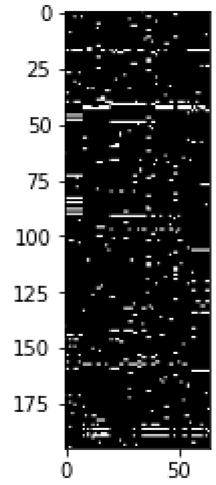


SUPPLEMENTARY TEXT FIGURE 2. Converted empirical dataset, with black pixels representing minor and white major alleles.

*ABC model selection*

In our ABC approach, the same strategy and prior values were used for simulation, but we performed 100,000 simulations and calculated Summary Statistics (SuSt) from the resulting datasets using the msSS.pl script (available at http://raven.wrrb.uaf.edu/~ntakebay/teaching/programming/coalsim/scripts/msSS.pl). This script calculates the following SuSt directly from the ms' output: pairwise differences between sequences ($\pi$); Tajima's D; Fay and Wu's $\theta_H$; the difference between $\theta_H$ and π (*H*); and π within and between every pair of lineages. We reduced the dimensionality of the SuSt with PCA, retaining the minimum number of axes that contained at least 99% of the total variance (as suggested by Perez et al. 2016). Cross-validation runs were performed with 100 pseudo-observed datasets to evaluate the capacity of our ABC implementation to predict the correct simulated model with a varying number of simulations. The empirical data were then used as input to perform a rejection step retaining only the 20% more similar simulations to approximate the posterior using the neural network algorithm implemented in the R package 'abc' (Csilléry et al. 2012).

*Drosophila dataset*

For the *Drosophila* datasets from Campillo et al. (2020), we also converted the empirical data for the *ms* format. To avoid discarding too many segregating sites due to missing data, we selected only loci with at least three samples from each species in the pair *D. melanogaster* - *D. simulans* (total of 625 segregating sites in *adh*, *amy* and *coII*) and three samples for *D. melanogaster* and two from *D. sechellia* (total of 185 segregating sites in *adh*, *amy* and *amyrel* for the species pair *D. melanogaster* - *D. sechellia*). We tested two concurrent scenarios: (1) all specimens assigned to only one species and (2) two separate species, in agreement with BPP and reproductive isolation metrics presented by Campillo et al. (2020). For both scenarios, we used a uniform distribution between 5 and 300 for $\theta$ and divergence times for the model with two species from a uniform distribution between 0.01 and 0.5 coalescent units. We evaluated the performance and predicted the most likely scenario for the empirical data using the same procedures described above for CNN and ABC in *P. aurisetus.* All scripts and datasets are available at

https://github.com/manolofperez/CNN_spDelimitation_Piloso.

**Supplementary Figures and Tables**

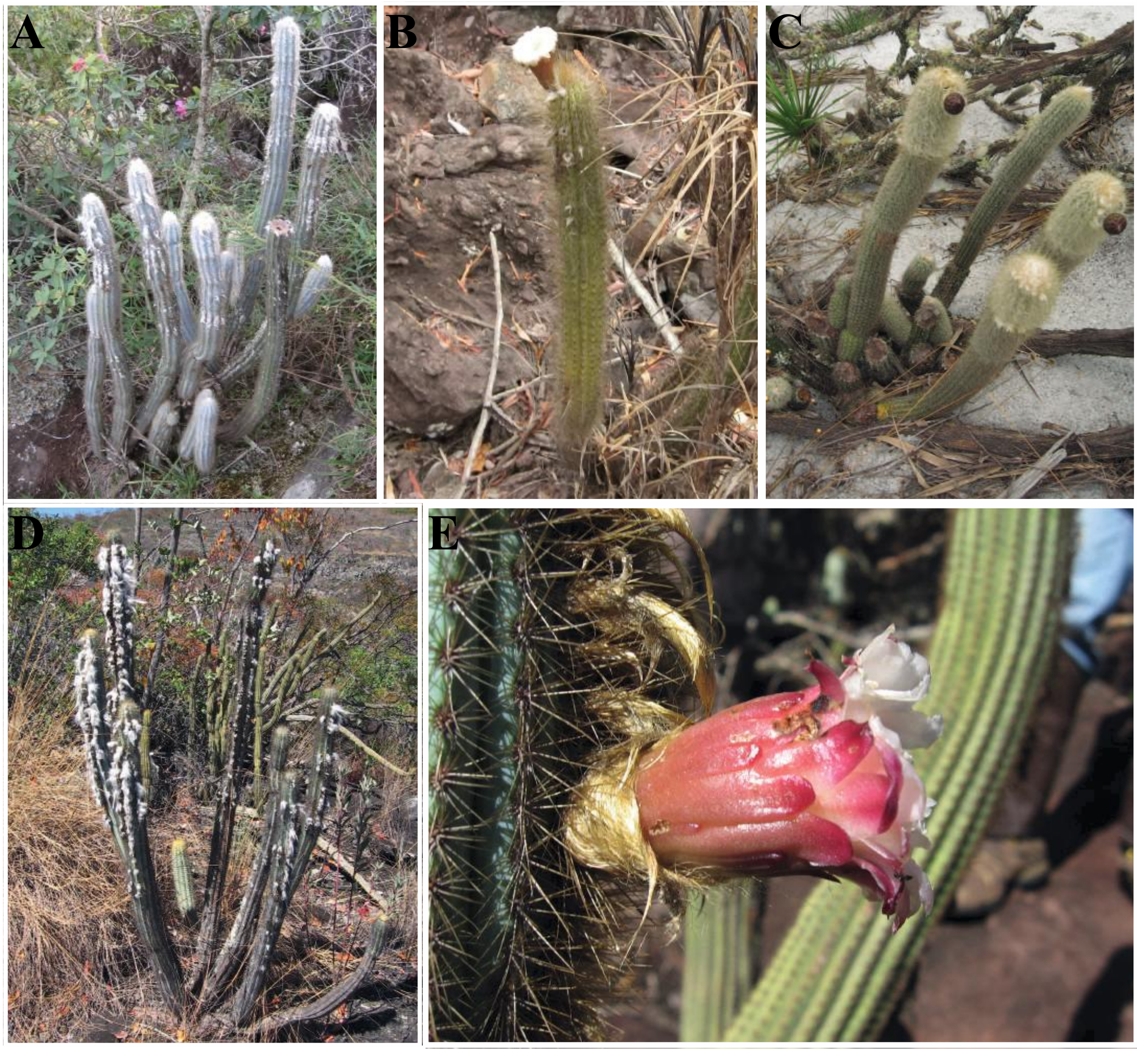


SUPPLEMENTARY FIGURE S1. Representatives of *P. aurisetus* subsp. *aurisetus* (A-D), and *P. aurisetus* subsp. *aurilanatus* (E) sampled in this study. A. *P. aurisetus* subsp. *aurisetus*; B. syn. *P. aurisetus* subsp. *werdermannianus*; C. syn. *P. aurisetus* subsp. *densilanatus*; D. syn. *P. aurisetus* subsp. *supthutianus*; E. *P. aurisetus* subsp. *aurilanatus*. Photo credit: EMM.

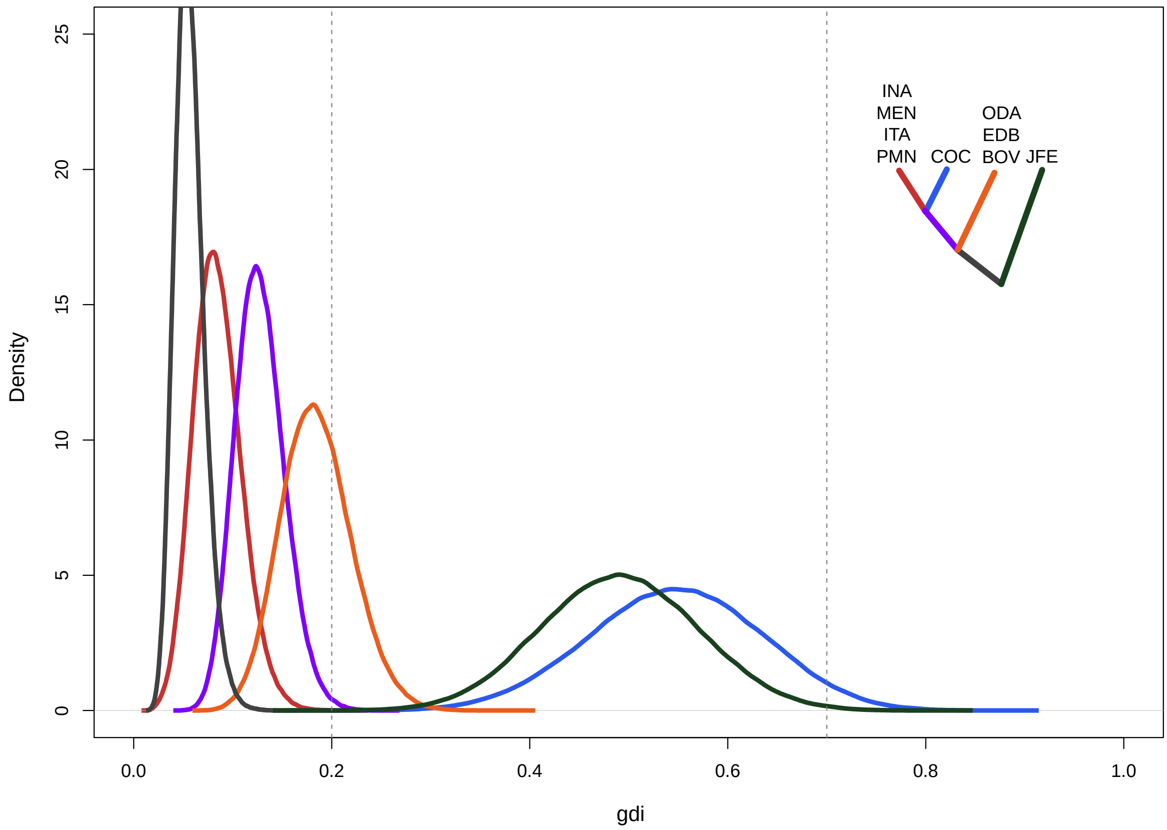


SUPPLEMENTARY FIGURE S2. Posterior distribution of genealogical divergence index (*gdi*) values generated by the BPP analysis. Color scheme according to branches from the tree depicted in the upper right corner, with locality names following Fig. 2.

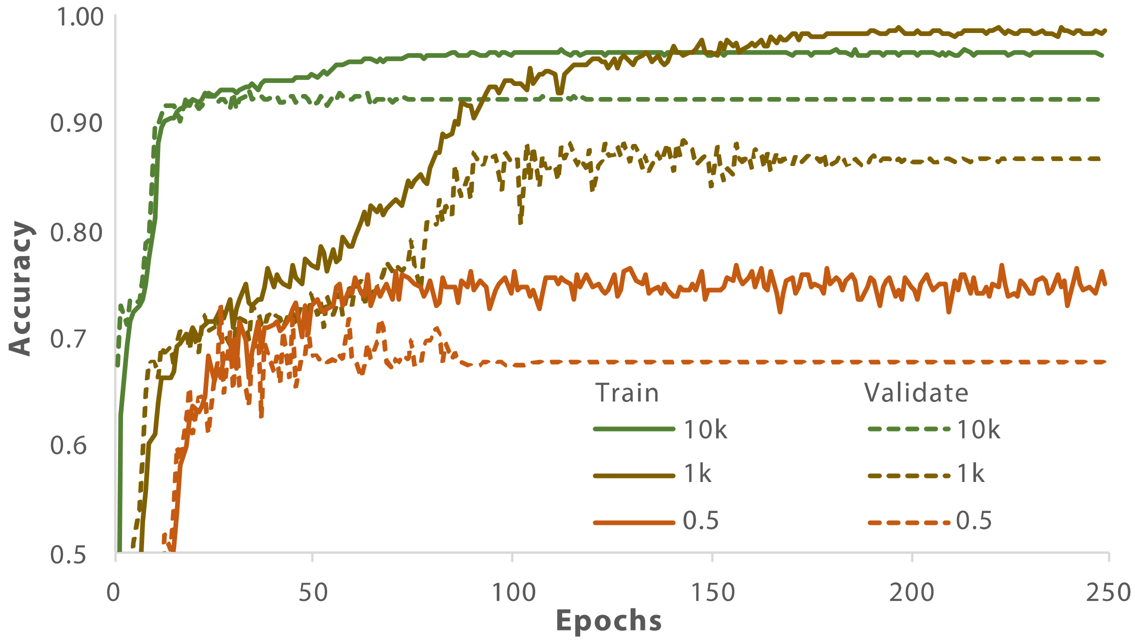


SUPPLEMENTARY FIGURE S3. Accuracy of the CNN along the training epochs with varying numbers of simulations (500 in orange, 1,000 in yellow and 10,000 in green) for the training (filled lines) and validation (dashed lines) sets.

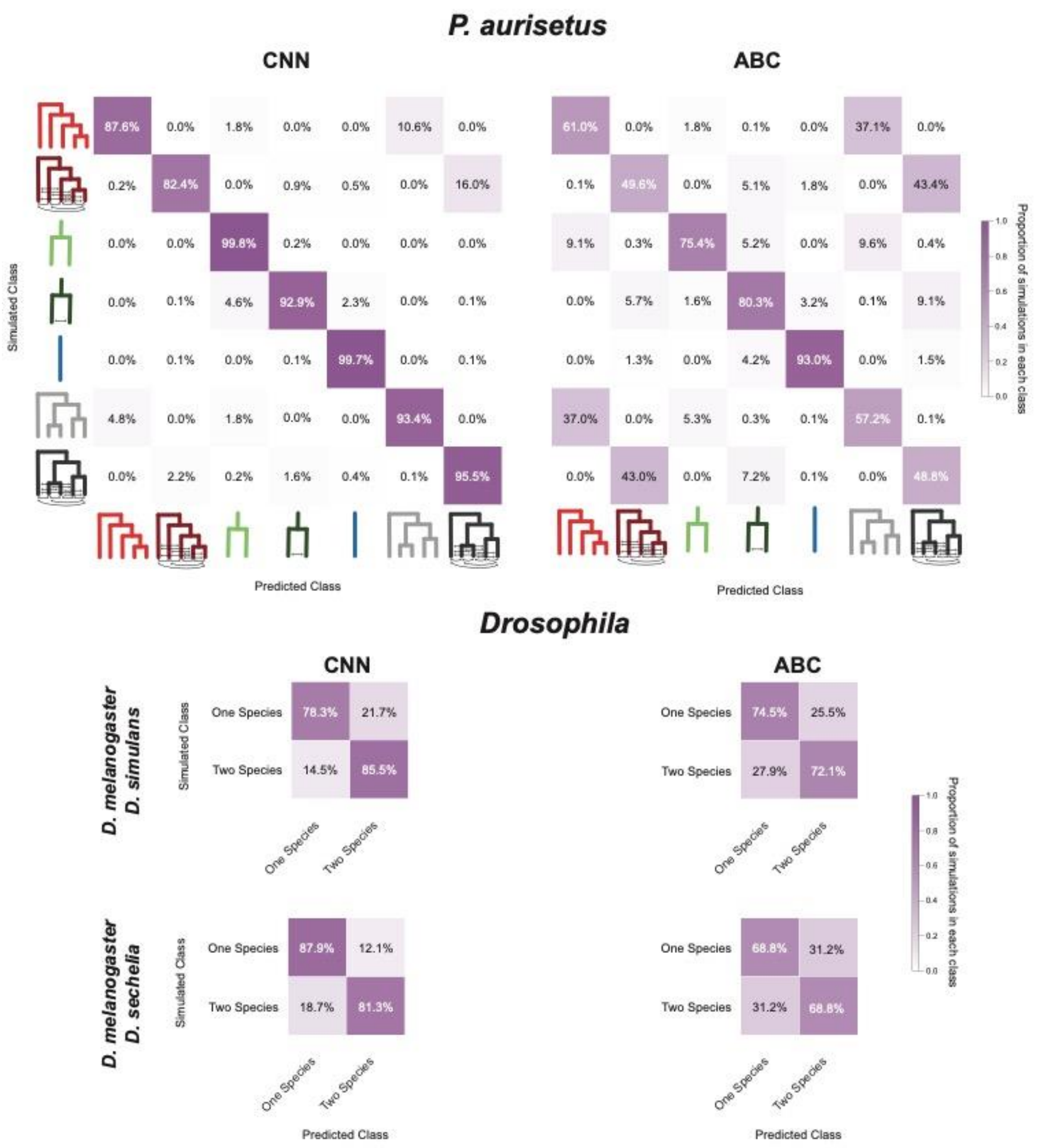


SUPPLEMENTARY FIGURE S4. Confusion matrices showing the percentage of predicted classes for each simulated scenario (model representation for *P. aurisetus* following Fig. 1) using test data (independent simulations not evaluated in the training step). The results are shown for both methods (CNN and ABC) in the three taxonomic systems analyzed (*P. aurisetus* and the species pais *D. melanogaster-D. simulans* and *D. melanogaster-D. sechellia*.

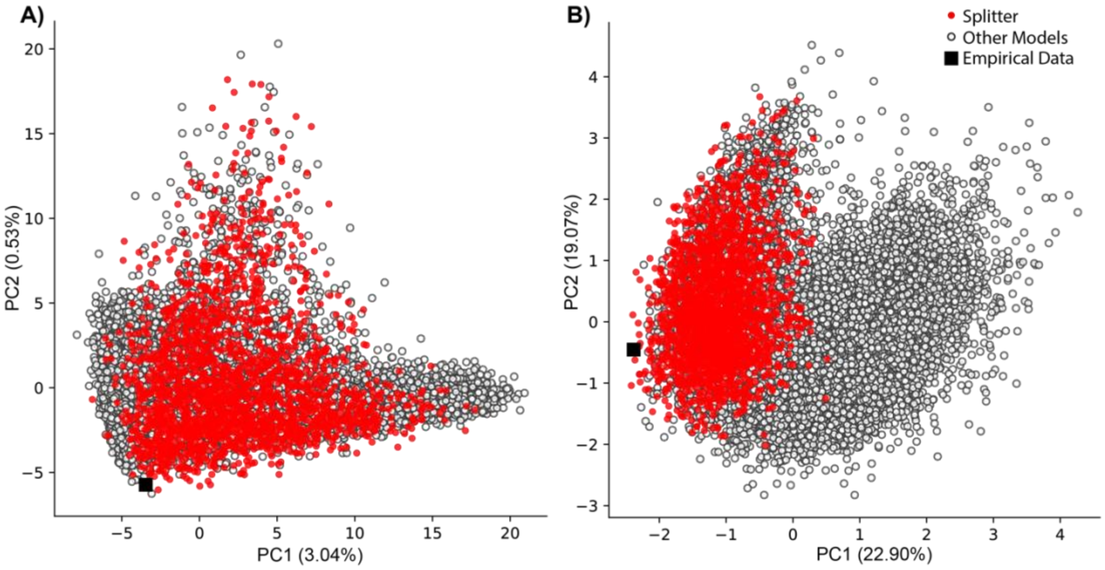


SUPPLEMENTARY FIGURE S5. Principal coordinate analysis for simulated (A -prior and B – ABC posterior) and empirical data in *P. aurisetus*. Simulations from the 'splitter' model are shown following the same color scheme in Fig. 1
