## Supplementary material for "Coalescent-based species delimitation meets deep learning: Insights from a highly fragmented cactus system": Table S1

Table S1. Primer sequence of each marker used in this study for microfluidic amplification. The genomic region for each marker is coded as follows: nr = nuclear, mt = mitochondrial, and cp = plastidial. Substitution models are provided for loci used in the final analyses

| Locus | Genomic Region | *Primer* Fwd (5´- 3´) | *Primer* Rev (5´- 3´) | Reference | Substitution Model |
|---|---|---|---|---|---|
| *PaANL*_8 | nr | TCCTCTCTTTTCTAGGGACGAC | CCCCATTCTTTCTTCATTCTATC | Perez et al. (2016b) | NA |
| *PaANL*_10 | nr | GAGAACGTCAATCCGACAGG | GAACATAGGCTGGCCTCTTC | Perez et al. (2016b) | NA |
| *PaANL*_15 | nr | GACCCTAACGAGGGTGAGAC | AAATCATTTCATGAGGCATCG | Perez et al. (2016b) | HKY+I |
| *PaANL*_17 | nr | TGTCCACCCCATAGAAGAGG | TTTAGATGAGTCCCAAAAGATACAC | Perez et al. (2016b) | K80+I+G |
| *PaANL*_28 | nr | CGTAGCAAACAGACATCCACTT | AAGAAATGCAACAAAAGAGTACCA | Perez et al. (2016b) | K80+I |
| *PaANL*_35 | nr | TCCTCTTTCCTACCATTCTTTCT | GTTTGAGGAAGGCAGAGGAG | Perez et al. (2016b) | K80+I |
| *PaANL*_46 | nr | ACTTTCCTGTRTCATATGTAA | CGAACTGGCCTCGGATTC | Perez et al. (2016b) | NA |
| *PaANL*_50 | nr | CGGGTCTAACTTGCCTTCAA | ACCCAACCGGTCAGATTGT | Perez et al. (2016b) | NA |
| *PaANL*_80 | nr | AAGAAGAACGGGCGAGTTG | AGGAGGTGGCAATGCAGTAG | Perez et al. (2016b) | K80+I |
| *PaANL*_82 | nr | CCAAGCAATATCGCATAAACAA | GGCACTAACTGATTCAATAACTGGT | Perez et al. (2016b) | NA |
| *PaANL*_87 | nr | TCTTTATGGCGTTATTCACTCG | CGAAGGCCTAACTTGACAGG | Perez et al. (2016b) | K80+I |
| *PaANL*_96 | nr | AGAAATGTGGGTCAGGAGGA | GAAATGCACATGCCTAGTGA | Perez et al. (2016b) | K80+I |
| *PaANL*_123 | nr | TTGCATGTTTATACAATTTTTCTTG | TGATAGATGCCAATCAGTCCAC | Perez et al. (2016b) | NA |
| *PaANL*_126 | nr | TCCTAAACAAGGGCTACGAAG | TGTACCAATGGGCAGCAC | Perez et al. (2016b) | TrN+I |
| *PaANL*_134 | nr | CGTGGTTTGACAAAACTTACCC | TCAGTGTTTCTAAGATGCTGCAC | Perez et al. (2016b) | NA |
| *PaANL*_140 | nr | TAGCCTCCTGAGCCCAAGC | GTTCATCAATGGGGAAGGTG | Perez et al. (2016b) | K80 |
| *PaANL*_142 | nr | CAAGCCTCTCCCTATAAC | TATAGAGTCTAGGCAAGGC | Perez et al. (2016b) | NA |
| *PaANL*_147 | nr | CTGTTGGCTCTGCATAGCTG | TGCTACACTGGCTTCATTGC | Perez et al. (2016b) | K80+I+G |

| | | | | | |
|---|---|---|---|---|---|
| *PaANL*_155 | nr | CTTTTCAGTCCAAAGCAAATTC | AAGGTCAGTAAGTCAAGCTCCTC | Perez et al. (2016b) | NA |
| *PaANL*_160 | nr | CGTGCTTTTACCTCCGTAAAG | CTAAGGGCTAATGGTGCTAGG | Perez et al. (2016b) | NA |
| *PaANL*_164 | nr | TTTCCCTTTATCCAAGTGACG | CGTGTTTGGTGGTCAATAGC | Perez et al. (2016b) | K80+I |
| *PaANL*_165 | nr | AGCCCTATATGTGGAAGG | GGAGTGCTTTCAAGCCTTTG | Perez et al. (2016b) | K80+G |
| *PaANL*_182 | nr | TTCAGGCTTAGGTTGGTGTTC | AGGGTCGTCACGATCATCC | Perez et al. (2016b) | K80+I |
| *PaANL*_187 | nr | CCGATTGAGGCTAGAAGCTG | TGTCTCTTGGCTTTACTTTAGGG | Perez et al. (2016b) | NA |
| *PaANL*_196 | nr | GCTTGGAGGTTTCCAATGAG | GAATGCTAAGGCCAAAAAGC | Perez et al. (2016b) | NA |
| *PaANL*_205 | nr | AAATCGGAGTCACAACAGAGA | TACCGAGATCTTGCGATGC | Perez et al. (2016b) | HKY+G |
| *its2* | nr | ATGCGATACTTGGTGTGAAT | GACGCTTCTCCAGACTACAAT | This Study | NA |
| *ppc* | nr | GAGATGAGGGCAGGGATGAGTTACTTCC | CTAGCCAACAAGCAAACATC | This Study | NA |
| *nhx1* | nr | TCTTCAGTGAAGATCTYTTCTTYAT | GAAGTTSCGGAARAATTGCTTCTT | This Study | NA |
| *isi1* | nr | AAGCKCCGAGTMTCSTGCTTT | AACTCYGCGCATTTCATCCT | This Study | NA |
| *apk1* | nr | GTTCCTGTTTCTGTTGTAGGTG | CAGCTTTGGAGTGGTTCTCC | This Study | K80+I |
| *cox1* | mt | CCAGTGAGTCCTCCAATGGT | TTTCGGTCATCCAGAGGTCTA | This Study | K80 |
| *cox2* | mt | TGATTCCTCACCTATCCTGTCA | CAGCGAGAAGCWAGGGATGT | This Study | K80 |
| *petA* | cp | TGCAACTAAAACAAGTTCTTGCTAA | TGGATTCACCCTCTGAAACA | This Study | NA |
| *psbA-psbB* | cp | ATCAGGAAGAACCGCTTGTG | TACTCCTTCTATTTCTACCCATCCA | This Study | NA |
| *psbC* | cp | TGAAGCTAATGCCCTTCCTC | GTGAGTTTTACGGGCCCACT | This Study | NA |
| *ycf1* | cp | GAGAAGGAGGTAGCCGCATC | TGTGACCAATTAACCAACCAACA | This Study | NA |
| *atpB-rbcL* | cp | GAAGTAGTAGGATTGATTCTC | CATCATTATTGTATACTCTTTC | This Study | HKY+G |
| *petL-psbE* | cp | AGTAGAAAACCGAAATAACTAGTTA | TATCGAATACTGGTAATAATATCAGC | This Study | HKY+G |

| | | | | | |
|---|---|---|---|---|---|
| *rpS16* | cp | AAACGATGTGGTARAAAGCAAC | AACATCWATTGCAASGATTCGATA | This Study | NA |
| *rpL16* | cp | AGAAGCGATGGGAACGATGG | TTCTTCTCATCCGGCTCCTC | This Study | HKY+G |
